# Imperfect Linkage Disequilibrium Generates Phantom Epistasis (& Perils of Big Data)

**DOI:** 10.1101/388942

**Authors:** G. de los Campos, D. Sorensen, M. A. Toro

**Author notes:** Corresponding Author. 775 Woodlot Dr. (1311), East Lansing, MI, 48824.

## Abstract

The genetic architecture of complex human traits and diseases is affected by large number of possibly interacting genes, but detecting epistatic interactions can be challenging. In the last decade, several studies have alluded to problems that linkage disequilibrium can create when testing for epistatic interactions between DNA markers. However, these problems have not been formalized nor have their consequences been quantified in a precise manner. Here we use a conceptually simple three locus model involving a causal locus and two markers to show that imperfect LD can generate the illusion of epistasis, even when the underlying genetic architecture is purely additive. We describe necessary conditions for such *“phantom epistasis”* to emerge and quantify its relevance using simulations. Our empirical results demonstrate that phantom epistasis can be a very serious problem in GWAS studies (with rejection rates against the additive model greater than 0.2 for nominal p-values of 0.05, even when the model is purely additive). Some studies have sought to avoid this problem by only testing interactions between SNPs with R-sq. <0.1. We show that this threshold is not appropriate and demonstrate that the magnitude of the problem is even greater with large sample size. We conclude that caution must be exercised when interpreting GWAS results derived from very large data sets showing strong evidence in support of epistatic interactions between markers.

## Introduction

A big challenge in genetics is to understand how variation at the DNA sequences translates into phenotypic variation. Genome-wide-association (GWA) studies address part of this challenge by testing for the association between phenotype (or a disease indicator) with genotype, one locus at a time. In the last decade, many GWA studies were conducted; these studies have reported thousands of SNP's (single nucleotide polymorphism) associated to complex traits and diseases (http://www.ebi.ac.uk/gwas).

Recently, several studies in model organisms (e.g., Mackay 2014), humans (Strange, Ask, and Nielsen 2013) and agricultural species (e.g., Huang, Xu, and Cai 2014), have used genotype data linked to phenotypes to investigate the presence of epistatic interactions between loci. Cordell (2002, 2009) and Wei, Hermani, and Haley (2014) provide comprehensive reviews of the methods commonly used to detect epistatic interactions.

There are several issues associated with studies aimed at detecting interactions, including matters of scale, the importance of the contribution of epistasis at the level of the genotype effects or at the level of the genotypic variance (e.g., Hill, Goddard, and Visscher 2008) and how an interaction detected in a linear statistical model may be associated to biological pathways that underlie a complex trait (e.g., Wang, Elston, and Zhu 2010; Aschard 2016). The latter becomes particularly problematic when the markers used to assess associations between SNPs and phenotypes (or a disease indicator) are in imperfect linkage disequilibrium (LD) with the alleles at the causal loci (i.e., those responsible for inter-individual genetic differences in a trait or disease phenotype). Under those conditions, evidence supporting the existence of a non-null interaction between markers do not necessarily provide definite evidence of epistasis at causal loci. Indeed, when the SNPs used in association analyses are in imperfect LD with the alleles at causal loci, linear regression on SNPs may lead to unaccounted variance, or *missing heritability* (e.g., Manolio et al. 2009; de los Campos, Sorensen, and Gianola 2015). Furthermore, the unaccounted additive signal may be correlated with interaction contrasts, thus creating the “illusion” of epistasis even for traits that are purely additive.

Several authors have expressed concerns about the role that LD can have on the detection of epistasis (e.g., Wei, Hermani, and Haley 2014). However, these problems have not been quantified nor have they been given a precise mathematical treatment. In this study, we present a simple three locus model involving a causal (unobserved) locus and two markers that makes explicit how *phantom epistasis* may emerge even in systems that are strictly additive. We use this model to derive a set of conditions that are necessary for the occurrence of phantom epistasis, and quantify the magnitude of the problem using simulations based on real human genotypes from the UK-Biobank. Our results suggest that imperfect LD can lead to seriously inflated type-I error rates. We also show that the rate of detection of phantom epistatic interactions increases with sample size; this should be considered when testing for epistatic interactions using big data sets such as the ones that are becoming available.

## Materials and Methods

To study what factors may induce phantom epistasis we consider a simple model with three bi-allelic loci. One of them, denoted as *z_i_*, represents a causal locus (also referred as to the ‘quantitative trait locus', QTL) and has a direct effect on the expression of a phenotype *y_i_*. The other two loci, denoted as *x*_1*i*_ and *x*_2*i*_, are markers that are possibly in LD with the QTL but have no causal effect on *y_i_*. For SNPs, a standard practice is to code genotypes (*z_i_*, *x*_1*i*_, *x*_2*i*_) by counting at each of the loci the number of copies of a reference allele carried by the *i*^th^ individual. Here, to facilitate the presentation we assume that genotypic codes and phenotypes are expressed as deviations from their corresponding means; therefore *E*(*z_i_*) = *E*(*x*_1*i*_) = *E*(*x*_2*i*_) = *E*(*y_i_*) = 0. In this setting, a single-locus strictly additive model takes the form

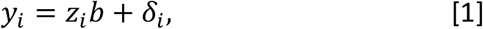

where *b* is the additive effect of an allele substitution at locus *z*, and *δ_t_* is an error term. Evidently, with only one causal locus there is no epistasis. We assume that [1] represents the causal model. Next, suppose that an instrumental regression of the form

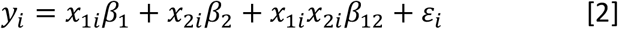

is used to investigate the presence of epistasis. Here, the *β's* are regression coefficients that are functions of the QTL effect (*b*) and of the (multi-locus) LD involving the two markers and the QTL genotypes. In the population, given the centered genotype codes, the regression coefficients entering in the right-hand-side of [2] are

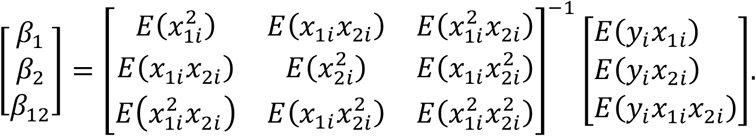

If the random residual *δ_i_* in expression [1] is orthogonal to the genotypes, then *E*(*y_i_x*_1*i*_) = *E*(*z_i_x*_1*i*_)*b*, *E*(*y_i_x*_2*i*_) = *E*(*z_i_x*_2*i*_)*b* and *E*(*y_i_x*_1*i*_*x*_2*i*_) = *E*(*z_i_x*_1*i*_*x*_2*i*_)*b*. Thus, the population regression coefficients are defined by

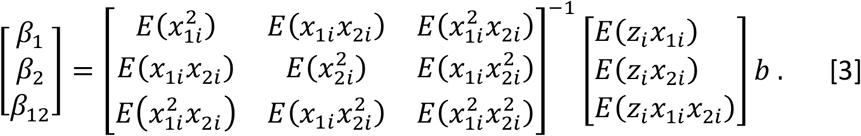

This indicates that the regression coefficients of the instrumental model [2] are not only functions of the QTL effect (*b*) and of pair-wise (1^st^ order) LD but also of higher order LD, e.g., joint disequilibrium at three loci, *E*(*z_i_x*_1*i*_*x*_2*i*_). The moments involved in the right hand-side of [3] are diploid genotypic measurements of LD. Under random mating these genotypic measures of LD are equal to twice the standard haploid measures of LD (the D-coefficients for two and tree loci linkage disequilibrium; see Section 1 of the Supplementary Methods for further details). In the population, the interaction effect *β*_12_ is given by a linear combination of two-loci LD between each of the markers and the QTL and by three-loci LD involving the two markers and the QTL: 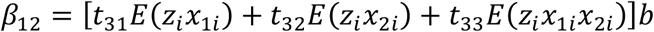. Here, the *t's* are the entries of the third row of the inverse of the coefficient matrix

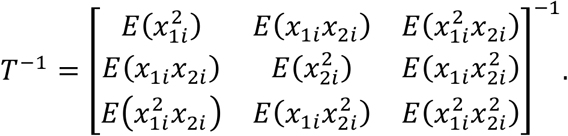

expressions to study the conditions that lead to a null interaction between markers.

### Conditions that lead to phantom epistasis

Next, we describe sufficient conditions for *β*_12_ = 0. These sufficient conditions also imply necessary conditions for phantom epistasis, *β*_12_ ≠ 0, to emerge.

#### Complete Linkage Equilibrium

If the QTL is in LE with the two markers, then (*z_i_*, *x*_1*i*_, *x*_2*i*_) = *p*(*z_i_*)*p*(*x*_1*i*_, *x*_2*i*_). Consequently, *E*(*z_i_x*_1*i*_ = *E*(*x*_1*i*_)*E*(*z_i_*) = 0, *E*(*z_i_x*_2*i*_ = *E*(*x*_2*i*_)*E*(*z_i_*) = 0, and *E*(*z_i_x*_1*i*_*x*_2*i*_) = *E*(*x*_1*i*_*x*_2*i*_)*E*(*z_i_*) = 0. Therefore, all elements of the right-hand-side of [3] are equal to zero and, thus *β*_1_ = *β*_2_ = *β*_12_ = 0. Therefore, *<u>a first necessary condition</u> for phantom epistasis to emerge is that the QTL must be in LD with at least one of the SNPs*.

#### Perfect Linkage Disequilibrium

On the other extreme, if there is perfect LD between the QTL and the marker pair (*x*_1*i*_*x*_2*i*_), then the QTL genotype can be expressed as a linear function of the two marker genotypes *z_i_* = *x*_1*i*_*β*_1_ + *x*_2*i*_*β*_2_. In this case, a linear regression on the two markers captures fully the QTL variance and therefore the interaction term will be equal to zero. (A derivation of this intuitive result is presented section 2 of the Supplementary Methods.) Therefore, perfect LD is a sufficient condition for *β*_12_ = 0. Consequently, *a <u>second necessary condition</u> for phantom epistasis to emerge <u>is imperfect LD</u> between the QTL and the marker pair*. This guarantees that some fraction of the QTL variance is not captured by linear regression on the two marker genotypes. Furthermore, if the left-out QTL signal is not orthogonal to the interaction contrast *x*_1*i*_*x*_2*i*_, then *β*_12_ ≠ 0.

#### Independence of one of the markers prevents phantom epistasis

Consider now an intermediate case where one of the markers (say *x*_2*i*_) is independent of the pair formed by the QTL and the other marker (*z_i_*, *x*_1*i*_). This implies that *p*(*z_i_*, *x*_1*i*_, *x*_2*i*_) = *p*(*z_i_*, *x*_1*i*_)*p*(*x*_2*i*_). Under this condition, because the two markers are in LE, the coefficient matrix and its inverse (*T*^−1^) is diagonal; therefore, 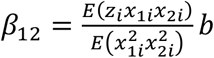. Moreover, *E*(*z_i_x*_1*i*_*x*_2*i*_) = *E*(*z_i_x*_1*i*_)*E*(*x*_2*i*_) = 0, implying that *β*_12_ = 0. Therefore, a *<u>third necessary condition</u> for phantom epistasis to emerge is that the three loci must be jointly in LD*.

In summary, *<u>phantom epistasis can emerge if the three loci are in mutual but imperfect LD</u>*.

### Simulation

The analytical results presented in the previous section indicates that multi-locus LD plays an important role in determining whether phantom epistasis may emerge. To shed light on the nature and the magnitude of the problem we conducted Monte Carlo simulations to assess how LD among the three genotypes (*z_i_*, *x*_1*i*_, *x*_2*i*_) affects the rates at which *H*_0_:*β*_12_ = 0 is rejected. Data were generated according to an additive model with a single causal locus that had strictly additive gene action (as in expression [1]) and then analyzed using an instrumental model such as the one in [2]. In this setting, rejection of *H*_0_: *β*_12_ = 0 is indicative of phantom epistasis.

Simulations were based on real human genotypes of distantly related white Caucasian individuals from the ***UK-Biobank***, a cohort study consisting of about half a million participants aged between 40-69 years who were recruited in 2006-2010. The National Research Ethics Committee approved the study and informed consent was obtained from all participants. Study details are described elsewhere (Sudlow et al. 2015).

To avoid confounding due to population structure and long-range LD due to family relationships we focused on distantly related white Caucasian individuals. Therefore, we only considered subjects whose self-reported ethnicity was Caucasian and confirmed their genetic race/ethnicity using SNP-derived principal components. From these individuals, we identified ~270,000 subjects that have pairwise genomic relationships, 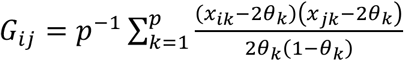, smaller than 0.03. Here, *x_ik_* and *x_jk_* are genotypes (coded as 0, 1, 2) at the *k^th^* SNP of the *i^th^* and *j^th^* individual, respectively, and *θ_k_* is the frequency of the allele counted at the *k^th^* loci. Genomic relationships were computed using the *getG()* function of the BGData R-package (Grueneberg and de los Campos 2017).

***Genotypes*** where from the Affymetrix UK BiLEVE Axiom and Affymetrix UK Biobank Axiom^®^ arrays. Only SNPs with minor-allele-frequency greater than 0.1% and those with missing calling rate smaller than 3% were used for simulations. Furthermore, since we focused on a single locus model, we used only SNPs mapped to chromosome 1. There were 66,331 SNPs mapped to chromosome 1, of those, 45,866 passed our minor-allele frequency and calling rate inclusion criteria.

#### Marker-QTL pairs

The position of the QTL genotype *z_i_* was determined by randomly choosing a marker position on chromosome 1. In a first simulation scenario, the two chosen markers were those flanking the QTL (i.e., those immediately adjacent to it). In subsequent simulation scenarios, the marker locus “to the right” (*x*_2*i*_) of the QTL *z_i_* was placed at increasing base-pair lags from the QTL, whereas the marker locus to the left (*x*_1*i*_) of *z_i_* remained always the most proximal marker “to the left of *z_i_*”. In this manner, the LD between one of the markers and the QTL was approximately constant whereas the LD between the distal marker, *x*_2*i*_, and the marker-QTL pair (*x*_1*i*_, *z_i_*) decreased as base-pair distance between the two markers increased. For each simulation scenario, we conducted 10,000 Monte Carlo replicates with random assignment of the QTL position within chromosome 1.

***Phenotypes*** were generated according to a single-locus additive model (expression [1]). with the QTL explaining one-half-of-one percent (0.005) of the phenotypic variance.

***Inferences*** were based on a linear model such as that of expression [2] extended with inclusion of an intercept and the top 5 SNP-derived PCs, that is

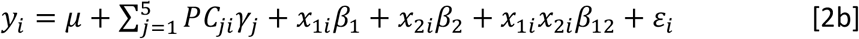

Principal components were included to avoid any confounding that may emerge from any remaining substructure that may have been present. The PCs used in [2b] were derived using 50K SNPs evenly distributed in the entire genome. The model of expression [2b] was fitted via least squares using the `ls.fit()` function of R (R Development Core Team 2012). Then for each scenario and MC replicate we saved the p-value associated to the interaction term and counted the proportion of times that *H*0: *β*_12_ = 0 was rejected when using a significance level of 0.05.

***Genotypic measures of LD*** between pairs of loci, *R*^2^(*x*_1_, *x*_2_), *R*^2^(*x*_1_, *z*) and *R*^2^(*x*_2_, *z*), were computed using the squared correlation between genotypes at the two loci. This information was stored for each MC replicate of each simulated scenario. The proportion of variance of the QTL genotype explained by linear regression on the two markers, *R*^2^(*z*~ *x*_1_ + *x*_2_), was computed by Analysis of Variance, of a linear model where the QTL genotype was regressed, via least squares, on the two markers using a main effects additive model of the form: *z_i_* = *μ* + *x*_1*i*_*a*_1_ + *x*_2*i*_*a*_2_ + *ε_i_*. The R-squared from this model was also saved for each MC replicate of each scenario and then used to analyze the relationship between this LD measure and the rate of rejection of *H*0: *β*_12_ = 0.

#### Effect of sample size

The power to detect a non-null interaction effect depends on two main factors: the proportion of variance explained by that interaction and sample size. The first factor is controlled in our simulation by controlling the distance between the QTL and the distal marker; this affects LD among the three loci and thus the size of the marker-interaction (see Methods). To assess the effect of sample size on inferences we carried out simulations using four different sample sizes: n=10K (K=1,000, this is representative of the size of a standard GWAS cohort) and n=50K, 100K and 250K (these sample sizes are more representative of modern large biomedical data sets).

### Data availability

The genotypes used in the simulation were from the UK Biobank. Data was acquired under project identification number 15326. The data are available for all bonafide researchers and can be obtained by applying at http://www.ukbiobank.ac.uk/register-apply/. The Institutional Review Board (IRB) of Michigan State University has approved this research with the IRB number 15-745.

## Results

**Figure 1** shows measures of linkage disequilibrium between the three loci (*z_i_*, *x*_1*i*_, *x*_2*i*_) involved in the system. The average (across Monte Carlo replicates) proportion of variance of the QTL (*z_i_*) explained by the most adjacent marker (*x*_1*i*_) averaged was about 0.085 (**Figure 1**); however, the distribution of this statistic is highly skewed. When *x*_1*i*_ and *x*_2*i*_ were the two flanking markers of the QTL, on average they jointly explained on average 15% of the QTL variance. Therefore, on average there was a sizable fraction of imperfect LD between the QTL and the markers. This leads to a sizable rate of “missing” heritability. The LD between *x*_2*i*_ and the pair (*x*_1*i*_, *z_i_*) decreased as the distance between *x*_2*i*_ and the pair (*x*_1*i*_, *z_i_*) increased. The R-sq. between *x*_2*i*_ and either the other marker or the QTL, falls very quickly for lags between 0-0.5Mb and reached near zero values at approximately 1 Mb (**Figure 1**).

**Figure 1.**
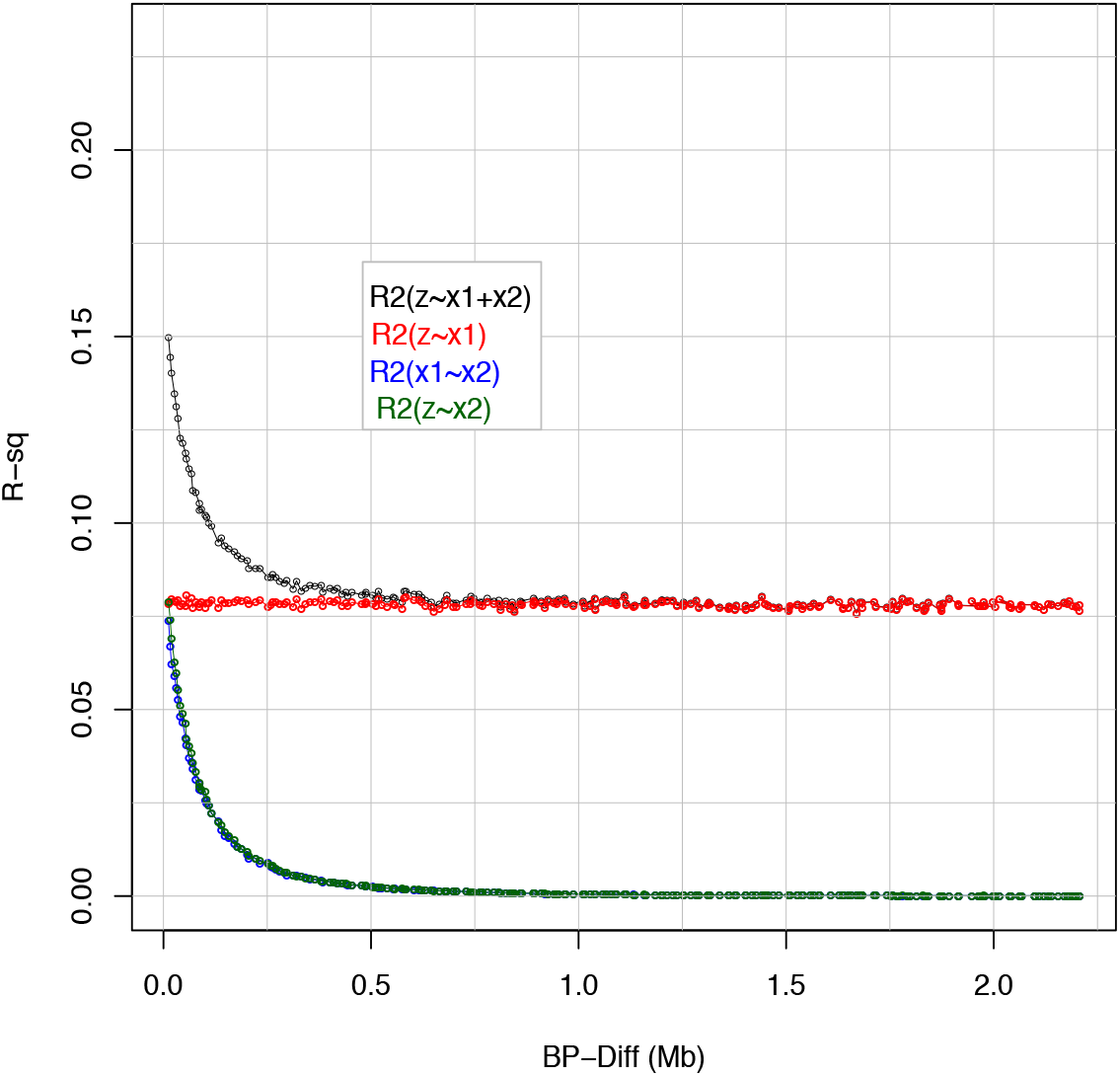
Average R-squared between pairs of loci and proportion of variance of the QTL genotype explained by the two markers, *R*^2^(*z_i_*~*x*_1*i*_ + *x*_2*i*_), versus distance between the QTL (*z_i_*) and the distal marker (*x*_2*i*_). Marker *x*_1*i*_ was always adjacent to the QTL.

In our simulation rejection of *H*_0_: *β*_12_ = 0 was performed using a significance level of 0.05. **Figure 2** displays the estimated rates of rejection by BP-distance between the QTL and the distal marker (*x*_2*i*_) and sample size. For the largest sample size, the curve relating empirical rejection rates with BP distance was clearly above 0.05 for distances of up to 2MB. The highest rejection rate was observed for *n*=250,000 when *x*_2*i*_ and the QTL were at a distance of about 0.15 MB; here the empirical rejection rate was ~0.13–this is more than twice the value expected under the absence of phantom epistasis. The curves relating empirical rejection rates with physical distance reach the nominal rejection rate of 0.05 at ~1Mb for n=10,000; however, for larger sample size the curves stayed above 0.05 even for distances longer than 1Mb.

**Figure 2.**
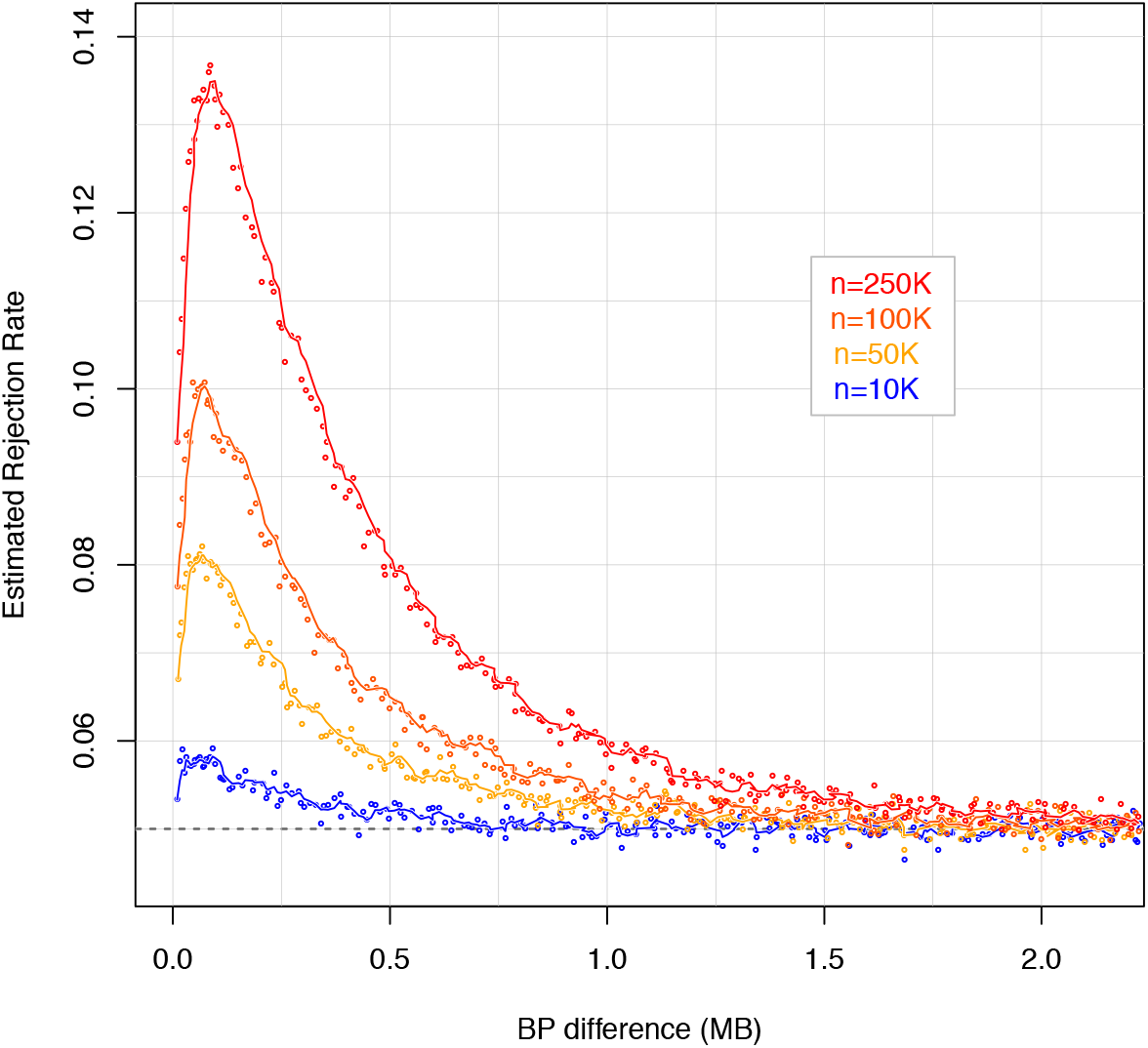
Estimated rejection rates versus distance between the QTL and the distal marker, by sample size. In the simulations one of the markers (*x*_1*i*_) was adjacent to the QTL (*z_i_*) and the other marker (*x*_2*i*_) was placed at increasing distance from the pair (*x*_1*i*_, *z_i_*).

The extent of LD varies substantially along the genome; therefore, for a given BP distance some regions may have very weak LD while others may have, at the same distance, SNPs in moderate or high LD. **Figure 3** shows another way of viewing the simulation results displayed in **Figure 2** where the average rejection rate is calculated within bins of R-sq. between the two markers. When the two markers were un-correlated, rejection rates were very close to 0.05 indicating absence of phantom epistasis. However very small LD between the two markers generates considerably higher rejection rates: an *R*^2^(*x*_1*i*_, *x*_2*i*_)~0.1 leads to rejection rates as high as 0.21 with the largest sample size. The maximum rejection rates occur when the R-sq. between markers is between 0.1 to 0.2. Beyond this value in the range (0.2-0.9) rejection rates shows a linear decline.

**Figure 3.**
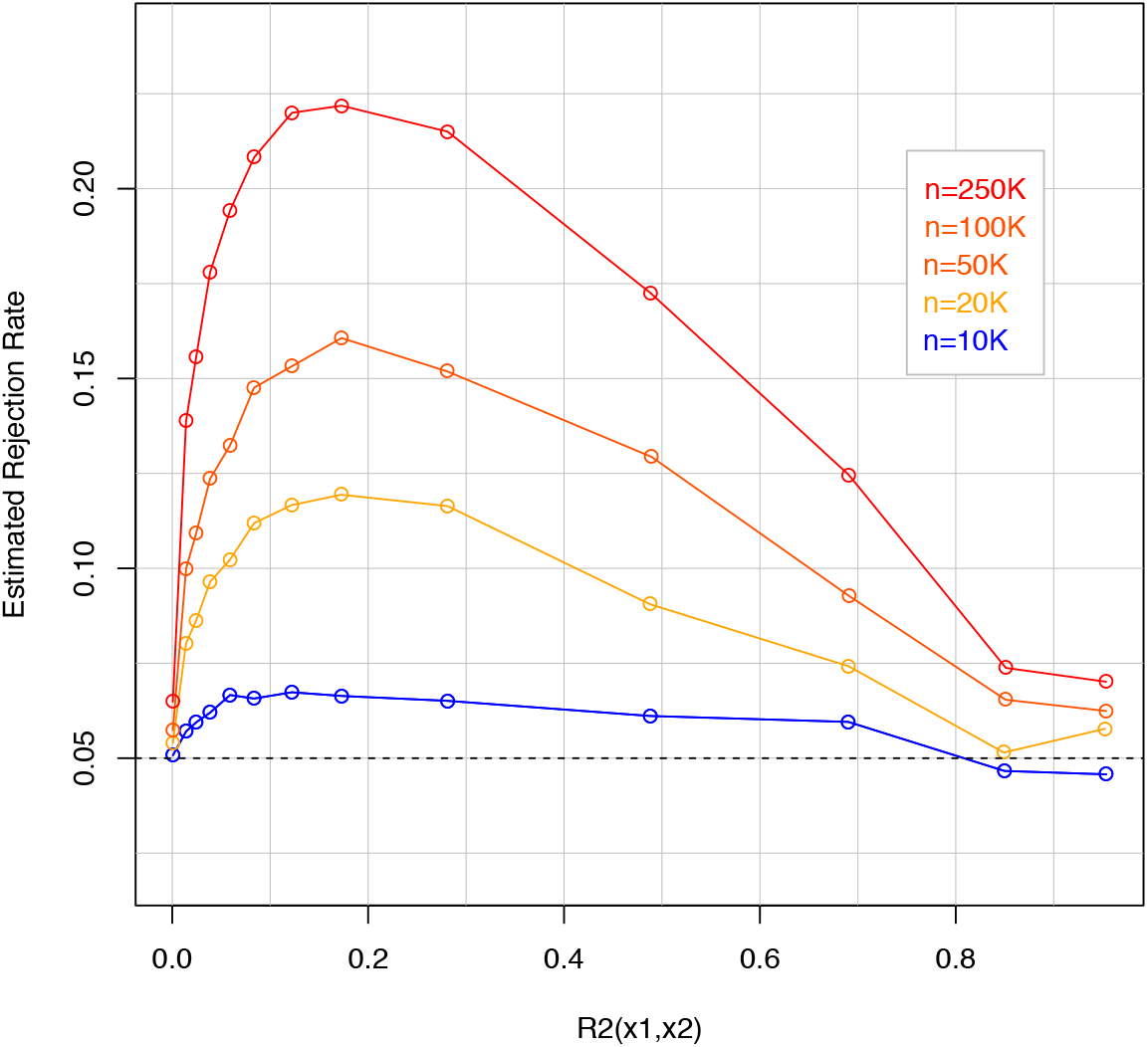
Estimated rejection rates by R-sq. between the two markers and sample size. In the simulations one of the markers (*x*_1*i*_) was adjacent to the QTL (*z_i_*)) and the other marker (*x*_2*i*_) was placed at increasing distance from the pair (*z_i_*, *x*_1*i*_).

The results in **Figures 2** and 3 are in line with the conceptual model described in the previous section. Analytically, the conditions needed for phantom epistasis to emerge include simultaneous but imperfect LD between the three loci. When the distal marker becomes independent of the QTL-proximal-marker pair, there is no phantom epistasis. This happens at about 2MB (**Figure 2**) and requires the R-sq. between the two markers to be very close to zero (**Figure 3**). When the LD between the QTL and the marker pair is very high but imperfect (e.g. 0.9 < *R*^2^(*z*~*x*_1_ + *x*_2_) < 1), some phantom epistasis is generated. However, for those R-sq. values the amount of signal that is not captured by the linear regression on the two markers and that can be recaptured by an interaction term involving both is small. Therefore, a very large sample size is required to detect the phantom epistasis (compare the empirical rejection rates in **Figure 3** for R-sq. in the range 0.9-1).

## Discussion

There is a substantial amount of literature reporting the presence of epistasis affecting complex traits but results, when scrutinized, have been controversial. Sometimes the controversy spawns from the suspicion that epistatic interactions may be capturing additive signals that were missed by the baseline additive model used to test interactions. For instance, Hermani et al. (2014) identified 30 pairs of SNPs that interact influencing gene expression and that were replicated across two independent studies. In a subsequent study (Wood et al. 2014) replicated many of the interactions reported by Hermani et al.; however, in each case, using sequence data, a single third variant could explain all the apparent epistasis. This happened even after removal of all pairs of SNPs with *r*^2^ < 0.1 which was suggested by Wei, Hermani, and Haley (2014) to minimize confounding due to “haplotype effects”.

However, the problem of why and under what conditions additive effects may generate “epistatic signals” has not be formalized. In this work, we use a simple three locus model to reveal the conditions that lead to phantom epistasis. We show that phantom epistasis emerges in the presence of simultaneous but imperfect mutual LD between the three loci (the QTL and the two markers involved in the interaction). This conceptually simple three loci model can be extended to more complex settings (e.g., multiple QTL-marker trios) without affecting the underlying source of the principle: if additive QTL variance is imperfectly captured by linear regression on markers and the unexplained variation is not orthogonal to interaction contrasts, then phantom epistasis emerges.

***Testing interactions among weakly correlated SNPs only*** (e.g., considering only SNP-pairs with *r*^2^ < 0.1) *is not a solution*. Our simulations demonstrate that phantom epistasis can emerge even when the two markers involved in the interaction are very weakly correlated. R-squared values greater than 0.05 or even smaller generate strong evidence of phantom epistasis particularly when sample size is large (**Figure 3**).

### Inferences under imperfect LD

In a series of recent studies, we (de Los Campos et al. 2013; de los Campos, Sorensen, and Gianola 2015b; Gianola et al. 2015) and others (e.g., M E Goddard 2009) have studied the role of imperfect LD on related inferential problems, including missing heritability (i.e., in generating a gap between the trait heritability and the amount of variance that can be captured by a SNP set) and whether imperfect LD can lead to estimates of genomic correlations between traits that are different than the underlying genetic correlations (Gianola et al. 2015). In all these cases, imperfect LD generates inferential difficulties; therefore, phantom epistasis should be seen as one of many issues arising when the markers used for inferences are in imperfect LD with causal variants.

### Perils of Big Data

The power to detect a small non-null interaction between markers emerging from phantom epistasis increases with sample size. Our simulation results demonstrate this clearly: for pairwise R-sq. between markers of 0.1 there are clear signs of phantom epistasis; however, rejection rates are not highly elevated over the significance level when sample size was moderate (n=10k) because at that R-sq. the size of the interaction effect is small and therefore the power to detect such small interaction effect with moderate sample size is low. Big Data is a blessing for genomic analysis of complex traits; however, some problems cannot be addressed with larger sample size. Moreover, in some cases, large sample size can make an inferential problem even more problematic.

### Dominance can also contribute to phantom epistasis

The conceptual and empirical model used in the simulation was based on a purely additive genetic architecture. In the presence of dominance, the true single-locus model becomes 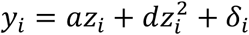 where *a* and *d* are additive and dominance values, respectively. If the empirical model of expression [2] is used to test for epistatic interactions then the left-hand-side of expression [3] remains unchanged, but the right-hand-side becomes

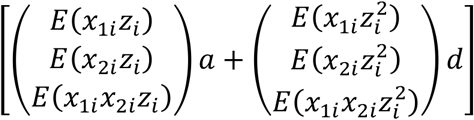

indicating that both dominance and additive effects can contribute to phantom epistasis. The conditions needed for phantom epistasis to emerge are similar to those under the pure additive model. These include, first, imperfect LD between 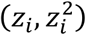 and the marker pair (*x*_1*i*_, *x*_2*i*_) such that neither *z_i_* nor 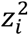 can be fully explained by a linear combination of the two markers. Secondly, phantom epistasis requires mutual LD at the three loci. If one of the markers (say *x*_2*i*_) is independent of the other-marker-QTL pair, then, 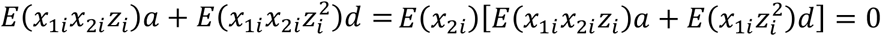.

***Local epistasis?*** Several studies have reported results highlighting the importance of ‘local’ epistatic interactions (e.g., Wei, Hermani, and Haley 2014; He et al. 2017). From a biological perspective it is plausible that multiple mutations in a gene may have collectively a larger impact than the simple sum of the effects of each mutation individually. And this could manifest as “haplotype effects” (e.g., Haig 2011). However, phantom epistasis provides an alternative explanation of why most of the epistatic interactions detected in GWAS occur between loci that are physically close. Indeed, we show analytically and empirically that LD between SNPs is required for phantom epistasis to appear, thus, phantom epistasis is expected to be predominantly a ‘local’ phenomena.

#### The additive-non-additive conundrum

Quantitative genetics studies properties of complex traits using regression analysis. In the field a careful distinction is made between observable and causal features of complex traits. For instance, it is well established that the linear regression of a phenotype on allele content yields estimates of the average effect of allele substitution and that both truly additive as well as dominance and epistatic effects can contribute to allele substitution effects. Furthermore, theoretical and empirical research has demonstrated that highly non-linear systems can generate signals that can often be explained almost completely with a linear model (Hill, Goddard, and Visscher 2008). For this reason, in general, one cannot make causal statements about gene action from observational variance component analyses (e.g., W. Huang and T. F. C. Mackay, 2016). Complicating matters even further we show in this study that the opposite can happen: under a purely additive model, imperfect LD can generate non-additive signals !

The recognition that phantom epistasis may be an important phenomenon does not negate the relevance of gene-gene interactions at the causal level. It simply stresses the difficulties that one faces when trying to learn about causal features of a system using observational data and inputs (markers) which are proxies for the underlying variants that may have causal effects on traits.

### Phantom epistasis: an opportunity to improve predictive performance?

In this work we have stressed that imperfect LD can limit the possibility to learn about causal effects. However, linear and non-linear genomic regressions can be very powerful predictive machines, and it is well-established that the model that is best for inferences is not necessarily the best predictive tool. Phantom epistasis creates inferential problems but also opens opportunities for improving prediction models. Indeed, by capturing signals that are missed by an additive model, non-linear models using interactions between markers may increase the amount of genetic variance captured and improve prediction accuracy. This may explain, for instance why some non-linear models such as kernel regressions have shown better predictive performance than additive models, especially in breeding populations with long-span LD and low marker density (de los Campos et al. 2010).

## Acknowledgments

The authors thank the participants and the personnel in charge of generating, curating and maintaining the data of the UK Biobank. GDLC and DS received financial support from NIH Grants R01GM099992 and R01GM101219. MAT received financial support from Spain's Ministry of *Economía y Competitividad* (grant CGL2016-75904-C2-2-P, MT). This work was supported in part by Michigan State University through computational resources provided by the Institute for Cyber-Enabled Research. We presented early versions of this work in a genetics seminar in George August Universitat and at the 2017 EAAP (European Federation of Animal Science) meetings. We are grateful for the comments received and would like to particularly thank the feedback offered by Johanes Martini.

## Supplementary Methods

### 1. Equivalence between genotype and haplotype measures of LD

In this section we present the standard haplotype two- and three-loci measures of LD and establish the connection between these measures and the genotype moments involved in expression [3].

#### Two-loci haplotype measure of LD

Consider a pair of bi-allelic loci (A and B, with alleles A_1_/A_2_ and B_1_/B_2_, respectively). The haplotype linkage disequilibrium parameter is *D_AB_* = *p*(*A*_1_, *B*_1_) − *p*(*A*_1_)*p*(*B*_1_). Let *X* = 1 when allele *A*_1_ is present and *X* = 0 when allele *A*_2_ is present. Likewise, let *Y* = 1 when allele *β*_1_ is present and *Y* = 0 when allele *β*_2_ is present. Then *E*(*X*) = *p*(*A*_1_), *E*(*F*) = *p*(*B*_1_), *E*(*A*_1_) = *p*(*A*_1_, *β*_1_) and the covariance between *X* and *Y* is *Cov*(*X, F*) = *E*(*A*_1_) − *E*(*X*)*E*(*F*) which reduces to *D_AB_*, thus

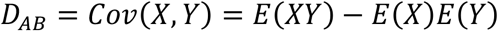

If the two genotypes are centered, then *E*(*X*) = *E*(*K*) = 0 and

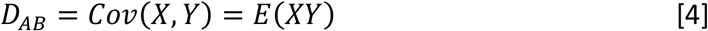

This is a haplotype analog of the 1^st^ order measures of LD entering in expression [3].

#### Three-loci haplotype measure of LD

For a system involving three bi-allelic loci (*A*, *B* and *C*, with alleles A_1_/A_2_, B_1_/B_2_, and C_1_/C_2_, respectively) a three-loci haplotype measure of LD can be defined as (Bennett 1954)

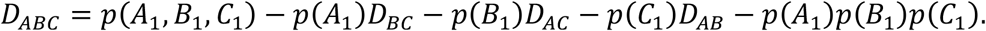

Extending the two-loci system by introduction of three binary random variables *X*, *F*, and *Z*, that take values 1 when the allelic forms *A*_1_, *B*_1_ and *C*_1_ are present, respectively, and take values 0 othewise, yields

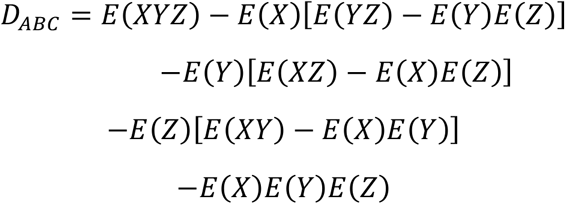

The three terms in square brackets represent pairwise disequilibria. When the three random variables are centered their marginal expectations are zero and the expression reduces to

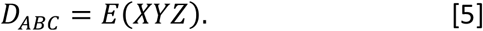

#### Relationship with genotype measures of LD

The disequilibria measures described above involve associations between alleles within gametes, whereas in the body of the paper, the expectations involve different genotypes. Assuming random mating, the expectations involving genotypes result in twice those involving gametes.

### 2. Perfect LD between markers and QTL prevents phantom epistasis

We demonstrate (the very intuitive result) that if a response (*z_i_*) can be fully captured by regression on a set of predictors (***x**_i_*), then the regression of *z_i_* on **x*_t_* plus ***w**_i_*,

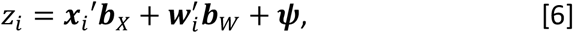

yields ***b**_w_* = 0 in the population.

#### Demonstration

In the population, the regression coefficients of [6], are defined by the following system

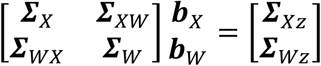

Where the ***Σ***'s represent covariance matrices: 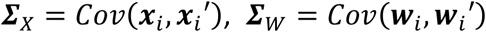 and 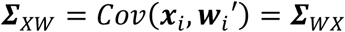. It follows that

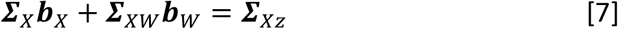

and

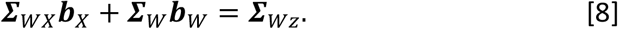

Solving [7] for ***b**_X_* yields 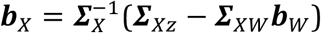. Plugging this into [8] yields, 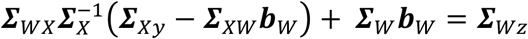. And solving for ***b**_W_* gives

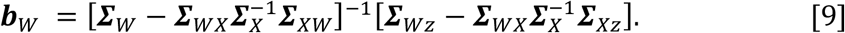

Now if *z_i_* can be fully explained by regression on ***x**_i_*, that is if *z_i_* = ***x**_i_′**a***, with 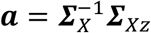, then, 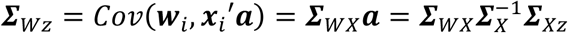, thus, 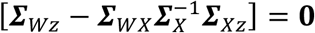 and therefore, ***b_W_*** = **0**. QED.

#### Implication

Let *z_i_* be the QTL genotype, ***x**_i_* = (*x*_1*i*_, *x*_2*i*_)′ be a vector containing the two marker genotypes and ***w**_i_* = *x*_1*i*_*x*_2*i*_ be the two-marker interaction contrast. Under perfect LD between the QTL and the markers the QTL genotype can be fully explained by linear regression on the two markers, that is *z_i_* = ***x**_i_′a*. Therefore, the above result (***b**_W_* = **0**) implies *β*_12_ = 0, i.e., absence of phantom epistasis.

## Literature Cited

Aschard, H. 2016. “A Perspective on Interaction Effects in Genetic Association Studies.” Genetic Epidemiology. https://doi.org/10.1002/gepi.21989.

Bennett, J H. 1954. “On the Theory of Random Mating.” Annals of Eugenics 18 (4):311–17. http://www.ncbi.nlm.nih.gov/pubmed/13148997.

Cordell, H J. 2002. “Epistasis: What It Means, What It Doesn't Mean, and Statistical Methods to Detect It in Humans.” Human Molecular Genetics 11:2463–68.

Cordell, H J. 2009. “Detecting Gene-Gene Interactions That Underlie Human Diseases.” Nature Reviews Genetics 10:392–404.

Gianola, D., G. de los Campos, M. A. Toro, H. Naya, C.-C. Schon, and D. Sorensen. 2015. “Do Molecular Markers Inform About Pleiotropy?” Genetics 201 (1):23–29. https://doi.org/10.1534/genetics.115.179978.

Goddard, M E. 2009. “Genomic Selection: Prediction of Accuracy and Maximisation of Long Term Response.” Genetica 136:245–52.

Grueneberg, Alexander, and Gustavo de los Campos. 2017. “BGData: A Suite of R-Packages for Analysis of Big Genomic Data [v 1.0.0].” Comprehensive R Archive Network (CRAN). https://cran.r-project.org/web/packages/BGData/index.html.

Haig, D. 2011. “Does Heritability Hide in Epistasis between Linked SNPs?” European Journal of Human Genetics 19:123.

He, Sang, Jochen C. Reif, Viktor Korzun, Reiner Bothe, Erhard Ebmeyer, and Yong Jiang. 2017. “Genome-Wide Mapping and Prediction Suggests Presence of Local Epistasis in a Vast Elite Winter Wheat Populations Adapted to Central Europe.” Theoretical and Applied Genetics 130 (4). Springer Berlin Heidelberg:635–47. https://doi.org/10.1007/s00122-016-2840-x.

Hermani, G, K Shakhbazov, H J Westra, T Esko, A K Henders, A F McRae, J Yang, et al. 2014. “Detection and Replication of Epistasis Influencing Transcription in Humans.” Nature 508.

Hill, W G, M E Goddard, and P M Visscher. 2008. “Data and Theory Point to Mainly Additive Genetic Variance for Complex Traits.” In PLos Genetics.

Huang, A, S Xu, and X Cai. 2014. “Whole-Genome Quantitative Trait Locus Mapping Reveals Major Role of Epistasis on Yield of Rice.” Plos One.

los Campos, Gustavo de, Daniel Gianola, Guilherme J. M. Rosa, Kent A. Weigel, and José Crossa. 2010. “Semi-Parametric Genomic-Enabled Prediction of Genetic Values Using Reproducing Kernel Hilbert Spaces Methods.” Genetics Research 92 (04). Cambridge University Press:295–308. https://doi.org/10.1017/S0016672310000285.

los Campos, Gustavo de, Daniel Sorensen, and Daniel Gianola. 2015a. “Genomic Heritability: What Is It?” Edited by Gregory S. Barsh. PLOS Genetics 11 (5):e1005048. https://doi.org/10.1371/journal.pgen.1005048.

los Campos, Gustavo de, Daniel Sorensen, and Daniel Gianola. 2015b. “Genomic Heritability: What Is It?” Edited by Gregory S. Barsh. PLOS Genetics 11 (5):e1005048. https://doi.org/10.1371/journal.pgen.1005048.

Los Campos, Gustavo de, Ana I Vazquez, Rohan Fernando, Yann C Klimentidis, and Daniel Sorensen. 2013. “Prediction of Complex Human Traits Using the Genomic Best Linear Unbiased Predictor.” Edited by Michael E. Goddard. PLoS Genetics 9 (7):e1003608. https://doi.org/10.1371/journal.pgen.1003608.

Mackay, Trudy F C. 2014. “Epistasis and Quantitative Traits: Using Model Organisms to Study Gene-Gene Interactions.” Nature Reviews. Genetics 15 (1):22–33. https://doi.org/10.1038/nrg3627.

Manolio, T A, F S Collins, N J Cox, D B Goldstein, L A Hindorff, D J Hunter, M I McCarthy, E M Ramos, L R Cardon, and \textitet. al. 2009. “Finding the Missing Heritability of Complex Diseases.” Nature 461:747–53.

R Development Core Team. 2012. “R: A Language and Environment for Statistical Computing.” Vienna, Austria. http://www.r-project.org/.

Strange, T, B Ask, and B Nielsen. 2013. “Genetic Parameters of the Piglet Mortality Traits Stillborn, Weak at Birth, Starvation, Crushing, and Miscellaneous in Crossbred Pigs.” Journal of Animal Science 91:1562–69.

Sudlow, Cathie, John Gallacher, Naomi Allen, Valerie Beral, Paul Burton, John Danesh, Paul Downey, et al. 2015. “UK Biobank: An Open Access Resource for Identifying the Causes of a Wide Range of Complex Diseases of Middle and Old Age.” PLoS Medicine 12 (3). Public Library of Science:e1001779. https://doi.org/10.1371/journal.pmed.1001779.

Wang, X, R C Elston, and X Zhu. 2010. “The Meaning of Interaction.” Human Heredity 70:269–77.

Wei, W.-H., G Hermani, and C S Haley. 2014. “Detecting Epistasis in Human Complex Traits.” Nature Reviews Genetics 15:722–33.

Wood, R W, M A Tuke, M A Nalls, D G Hernandez, S Bandinelli, A B Singleton, D Melzer, L Ferrucci, T M Frayling, and M N Weedon. 2014. “Another Explanation for Apparent Epistasis.” Nature 508.

